# Division of labor, bet hedging, and the evolution of mixed biofilm investment strategies

**DOI:** 10.1101/107102

**Authors:** Nick Vallespir Lowery, Luke McNally, William C. Ratcliff, Sam P. Brown

**Author notes:** Current address: META Center for Systems Biology, University of Oregon, Eugene OR 97403, USA. Correspondence to: Nick Lowery or Sam Brown.

## Abstract

Bacterial cells, like many other organisms, face a tradeoff between longevity and fecundity. Planktonic cells are fast growing and fragile, while biofilm cells are often slower growing but stress resistant. Here we ask: why do bacterial lineages invest simultaneously in both fast and slow growing types? We develop a population dynamical model of lineage expansion across a patchy environment, and find that mixed investment is favored across a broad range of environmental conditions, even when transmission is entirely via biofilm cells. This mixed strategy is favored because of a division of labor, where exponentially dividing planktonic cells can act as an engine for the production of future biofilm cells, which grow more slowly. We use experimental evolution to test our predictions, and show that phenotypic heterogeneity is persistent even under selection for purely planktonic or purely biofilm transmission. Furthermore, simulations suggest that maintenance of a biofilm subpopulation serves as a cost-effective hedge against environmental uncertainty, which is also consistent with our experimental findings.

## INTRODUCTION

After billions of years of evolution, many organisms retain an impressive capacity for innovation and adaptation to their environment. However, core traits such as durability and reproductive rate are often optimized such that improvements in one will often come at the cost of another - indeed, understanding how adaptation occurs when key fitness parameters are constrained by tradeoffs lies at the core of life history theory (1). While most life history theory has been developed with large multicellular organisms in mind, microbes also exhibit classical trade-offs in fecundity and longevity, with faster growing lineages tending to be more fragile (2, 3). Understanding how microbes manage such trade-offs remains a major goal in microbiology, both from mechanistic (4) and ecological (5) perspectives.

Multiple mechanisms of enhancing durability and longevity are available to microbes, but typically come at the cost of reduced metabolic proficiency. Spore formation is perhaps the clearest example of a high survival, low fecundity phenotype: by encasing the genome and some essential metabolic machinery in a thick and extremely resistant cell wall, dormant spores can survive for extraordinarily long durations (6, 7). Alternatively, cells may form metabolically dormant persister cells capable of surviving diverse environmental insults (8, 9). Finally, many microbial species form biofilms, where dense cell packing and production of a protective extracellular matrix provides broad resistance to stressors such as desiccation, predation, or chemical insult (10), but also limits space and nutrient diffusion, thereby reducing growth rates.

Clonally reproducing microbes present an interesting and experimentally tractable system to examine mixed-behavioral strategies. Across many species of microbe, single genotypes can produce coexisting subpopulations of rapidly dividing planktonic cells and slow-growing or dormant stress-tolerant phenotypes, but focus is often given to a specific phenotype of interest rather than the balance between alternate phenotypes. In this study, we examine how the trade-off between survival and growth of individual cells drives the evolution of mixed biofilm / planktonic investments on a lineage scale under diverse environmental conditions. Specifically, we build population dynamical models of bacteria in patchy environments, where cells can switch between biofilm and planktonic states within ephemeral patches (via planktonic colonization of the biofilm, and dispersal from the biofilm to the planktonic state), and can also transmit among patches either as biofilm or planktonic cells. We then ask, under what conditions is investment into biofilm favored, given that biofilms grow more slowly? If only one phenotype (i.e. biofilm or plankton) is favored for transmission to a new patch, does it ever pay to diversify into the cell type that is, from a transmission perspective, a ‘dead end’? Our model predicts that phenotypic diversification can pay across a range of environmental conditions, as rapidly growing planktonic cells can function as a ‘growth engine’ providing higher levels of future planktonic and biofilm cells for transmission. We then test our model predictions using stochastic simulations and experimental evolution of biofilm allocation in the environmental microbe *Pseudomonas aeruginosa*.

## RESULTS

### Biofilm Growth Dynamics

While it is well known that planktonic cells accumulate exponentially in nutrient-rich environments, it is less clear whether close-packed biofilm cells would follow the same functional form (note that here we are considering biofilm growth in the absence of any coupling with the planktonic compartment, i.e. no cells dispersing from the biofilm or colonizing from the bulk). We hypothesize that sparse colonization allows for lineages to grow exponentially, as there is little steric inhibition or nutrient depletion to slow growth. However, once confluence across the surface is reached, further growth is restricted to a fixed depth within the outermost layer in biofilms due to space and diffusion limitations (11, 12). We explore this conjecture using the individual-based simulation platform iDynoMiCS (13) to simulate a simple two-dimensional biofilm, and find that after an initial period of exponential growth, cell accumulation decays to a linear function in time (Figure 1). More generally, we find that the rate of biofilm growth depends on the geometry of the system being considered (Supplemental Text 1), with the growth rate following a polynomial of order equal to the dimensionality of the system (i.e. for a three dimensional sphere, biofilm cells accumulate as a cubic in time). However, it should be noted that for finite volumes there is a constant downward pressure through time on the order of the growth polynomial as the biofilm reaches confluence across each dimension (e.g., initially cubic expansion in three dimensions will decay to quadratic expansion in two dimensions once the limit in the z-direction is reached). These findings highlight that while biofilm cells do not face the extreme growth penalty of resistant spores or persister cells, they face a significant and compounding growth deficit in comparison with the exponential growth of planktonic populations.

**Figure 1:**
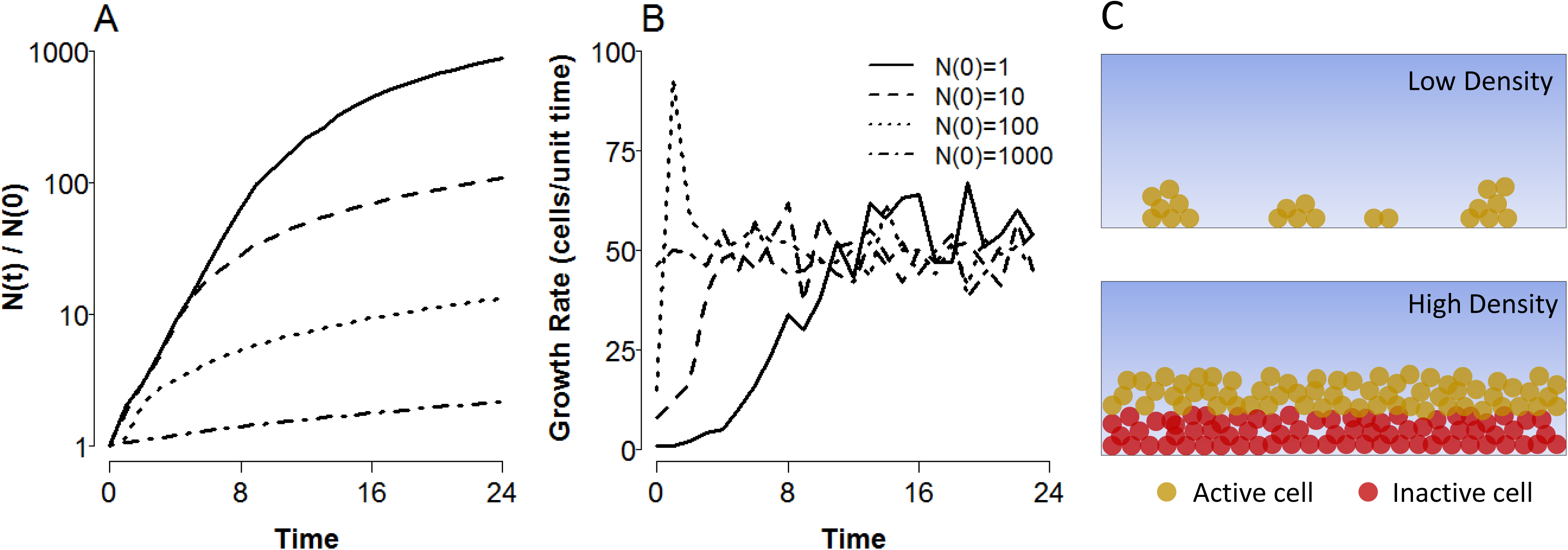
Biofilms grow sub-exponentially. **A**, accumulated cells and **B**, lineage growth rate through time for agent-based simulations of a 2-D biofilm growing on a 1-D surface. Varying inocula (see legend) were allowed to grow for a fixed time period under nutrient-rich conditions. Simulations were implemented using the agent-based simulation platform iDynoMiCS (13) **C**, Schematic depiction of nutrient depletion leading to growth arrest in the biofilm interior. Diffusion and active consumption of nutrients in the outermost layers of the biofilm (yellow cells) result in starvation conditions for cells in the interior regions (red cells).

### Coupled Biofilm-Plankton Dynamics

We next model a growing bacterial microcosm within which cells grow in one of two compartments, the planktonic (P) phase within the bulk fluid, and the biofilm (B) phase attached to a surface and in contact with the bulk. The compartments are coupled, such that biofilm cells can disperse to the planktonic phase, and planktonic cells can colonize the biofilm. Cells in each compartment also divide, with planktonic cells growing exponentially, and biofilm cells growing linearly (i.e. we assume a finite 2-D surface available for biofilm colonization, and ignore the initial super-linear growth period).

With the biofilm limited to linear expansion, we reasoned that the effects of growth within and dispersal from the biofilm would be negligible when coupled to exponential growth in the planktonic phase. This simplification was shown to be reasonable by comparing numerical simulations of the full model and a simplified model with no biofilm growth or dispersal (Figure S1), yielding the model system outlined in Figure 2A, and Equations 1.1. 1.2, 2.1 and 2.2 for within-patch growth. By setting the growth of the biofilm to zero, this simplified framework renders biofilm cells functionally equivalent to spores or persisters as described above, i.e. a subpopulation of non-dividing cells supported by the growth of vegetative cells, which presumably must provide some other benefit (e.g. environmental resistance) to the overall population to counteract this loss in fitness or else be lost from the population.

Our simplified model framework results in the following coupled differential equations:

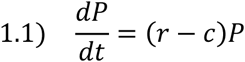

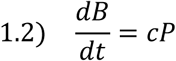

where *r* is the exponential growth rate, *c* is the rate of colonization of the biofilm (with 0 ≤ *c* < *r*). Note that in general *c* need not be bounded by *r*, and arbitrarily high values for *c* would result in a decline in *P* as switching to the biofilm phase outpaces planktonic growth, giving a sharp trade-off between the two compartments; we discuss this case in the context in which it arises below. Solving equations 1.1 and 1.2 as a function of time yields our within-patch population model,

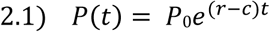

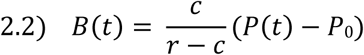

where *P_0_* is the planktonic inoculum.

The within-patch model reveals a temporal trade-off in biofilm accumulation with increasing colonization (Figure 2). As expected, planktonic cells decline monotonically with increasing colonization rate *c*, as more cells are siphoned from the planktonic to the biofilm compartment (Figure 2, A and C). The biofilm, however, shows more interesting dynamics with changing rates of colonization (Figure 2, B and D). At *c* = 0, no biofilm cells accumulate, and as *c* approaches *r*, all new planktonic cells colonize the biofilm, resulting in a static planktonic population and linear accumulation of biofilm cells (Figure 2A and B, yellow lines). However, when the planktonic fraction is allowed to expand exponentially (with 0 < *c* < *r*), the biofilm also accumulates cells roughly exponentially (once *P(t)* >> *P0*) at a constant fraction 
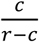
 of the planktonic population. High colonization rates thus provide more biofilm cells at short time scales, while lower colonization rates maximize biofilm over longer periods of growth (Figure 2B).

**Figure 2:**
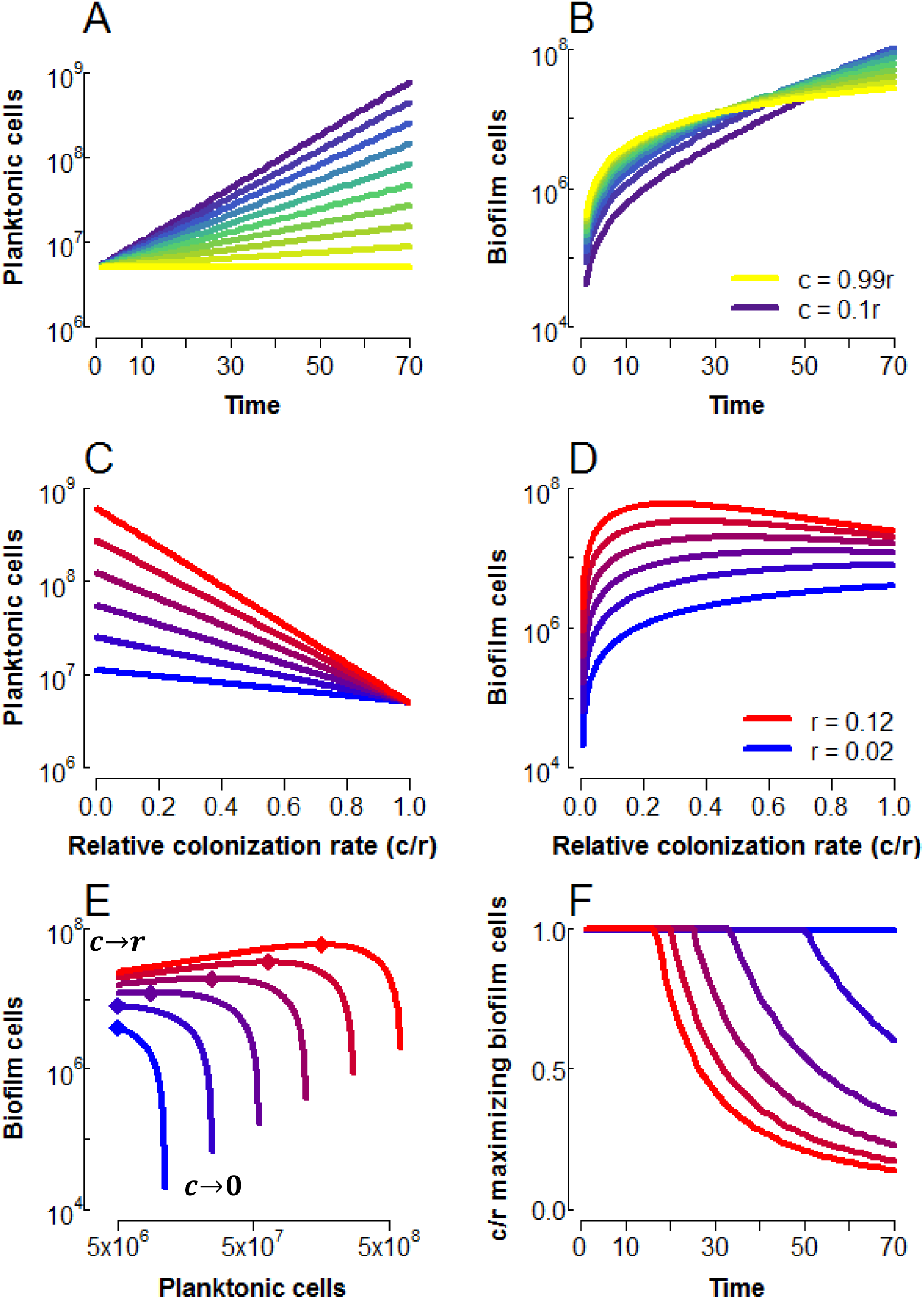
Maximal biofilm can be driven by planktonic growth. **A**, time series of planktonic and **B**,biofilm cells, for colonization rates *c* between 0.1*r* (purple) and 0.99*r* (yellow), with *r* = 0.08, *P0* = 5,000,000. **C**, planktonic, and **D**, biofilm cells as a function of colonization rate relative to growth rate for *t* = 40, 0.02 ≤ *r* ≤ 0.12 (blue-red color scale), *P0* = 5,000,000. **E**, Biofilm cells plotted against planktonic cells under the same conditions as C and D. The colonization rate varies over each curve, with the end points indicated by labels (relative colonization *c/r* approaches 1 at the left, and zero at the right). Diamonds indicate the maximum in biofilm cells. **F**, Relative colonization rate *c/r* at which biofilm is maximized. Note in panels D and E the limit of *c* = 0 is omitted, as this prevents any formation of biofilm cells

In Figure 2D, we find that the colonization rate maximizing biofilm cells declines with increasing planktonic growth rates, giving a humped shape in biofilm as a function of colonization rate. We can find an analytical condition for this relationship by examining the slope of *B* as a function of *c* as *c* approaches *r* (Equation 2.3, see Supplemental Text 2 for derivation).

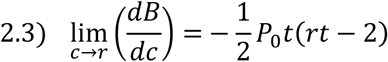

If the slope is negative, this would imply an interior maximum in *B* at some *c* < *r* (as biofilm cells necessarily increase as *c* increases from zero). We find that this limit is negative for *rt* > 2, i.e. the presence of a humped relationship in *B* requires patch quality (the product *rt*) to exceed a minimal threshold value.

These results suggest that the colonization rate maximizing biofilm will depend on opportunities for growth in the planktonic state (governed by growth rate *r* and growth duration *t*), which we explore further in Figure 2E and F. In Figure 2E, plotting biofilm cells against planktonic cells reveals that while limited growth (blue lines) leads to an allocation trade-off (i.e. increasing *B* necessarily comes at the cost of decreasing *P*), increasing growth rate decouples this trade-off, with *B* maximized at diminishing colonization rates *c*, depicted explicitly in Figure 2F. However, as colonization decreases beyond this point, biofilm declines sharply as colonization tends to zero.

Under high growth regimes, the planktonic fraction may therefore be viewed as a ‘growth engine’ to maximize biofilm: when within-patch conditions are sufficiently favorable to planktonic division (Equation 2.3), reducing the biofilm colonization rate *c* below the maximum increases the net flux of cells into the biofilm by expanding the pool of dividing planktonic cells, *P*. This growth engine effect is sufficient to drive colonization rates maximizing biofilm down to a fraction of the growth rate (Figure 2F).

### Evolutionary Model

While intermediate colonization rates may maximize biofilm, we note that any allocation towards the biofilm comes at the cost total population size in Equation 2.4:

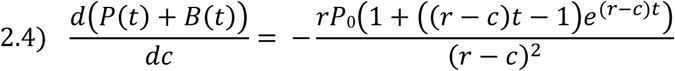

which is strictly negative for *t* > 0 and equal to 0 at *t* = 0. Given this trade-off between the biofilm and total population size, what conditions would favor biofilm investment (*c* > 0)? We can examine the evolutionary consequences of allocation in the within-patch population model by constructing a life cycle in which a population colonizes successive patches through space and/or time (i.e. migration between patches, or remaining in a single patch that experiences periodic disturbances). We define a fitness function (Equations 3.1 and 3.2) by assigning transmission probabilities *k_p_* and *k_b_* that a given cell from the respective planktonic or biofilm compartments will go on to found a new patch:

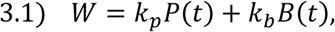

or, explicitly, per-founding cell (analogous to the reproductive number ‘R0’ framework common to parasite epidemiology and evolution, e.g. in (Frank, 1996))

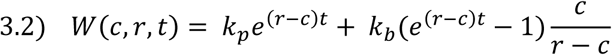

Equation 3.2 allows us to interrogate the fitness consequences of biofilm investment strategies (colonization rate *c*) across a wide array of ecological parameters. *k_p_* and *k_b_* capture the reproductive value of each cell type, and can be interpreted equivalently as a per-cell transmission probability or as the fraction of each subpopulation able to transmit successfully; they will dictate how well a given cell type (biofilm or planktonic) can survive the inter-patch transition, and be influenced by the nature of the environment. Growth time *t* describes the disturbance regime, i.e. how long a population can stay in a single patch, and *r* measures the nutrient quality of individual patches, or how rapidly planktonic cells can divide within the patch. We define *c*\* as the optimal colonization rate maximizing fitness *W* under a given ecological condition, and display the behavior of *c*\* as a function of transmission parameters *k_p_* and *k_b_* in Figure 3.

**Figure 3:**
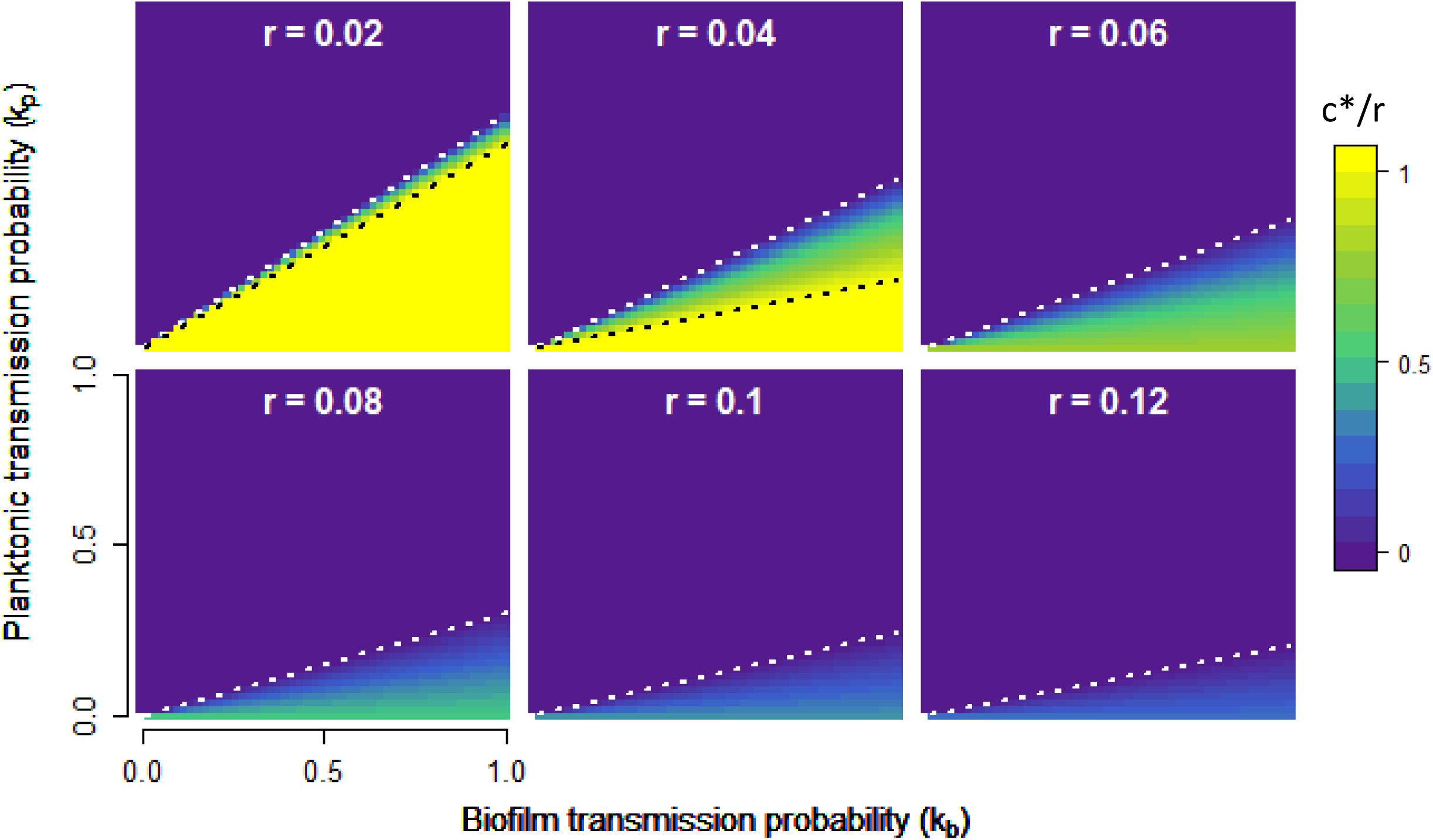
Optimal colonization rate *c*\*as a function of reproductive value for planktonic and biofilm cells. Contour plots showing the relative colonization rate (*c*/r*, yellow-purple color scale) optimizing fitness (Equation 3.2). Each panel displays *c*/r* as a function of *kp* and *kb*. Across panels, the growth rate *r* increases from *r* = 0.02 to 0.12, with *t* = 40 in all cases; similar plots varying *t* are displayed in Figure S2.White dotted line indicates the threshold at which *c*\* > 0, and black dotted line indicates the threshold at which *c*\* < 1 (see main text and Supplemental Text 3).

There are three general strategies microbes may adopt in maximizing fitness: devoting all resources to the biofilm fraction (*c*\* = *r*, below black dotted lines in Figure 3), splitting resources between the two fractions (0 < *c*\* < *r*, between dotted lines in Figure 3), or devoting all resources to the plankton (*c*\* = 0, above white dotted lines in Figure 3). In Supplemental Text 3, we investigate the conditions governing the two transitions defining these regimes by examining the behavior of Equation 3.2 in more detail.

### Trade-offs in allocation

When growth opportunities are limited (Figure 3, upper left panel) we see evidence of a sharp tradeoff between *B* and *P* investment, with very little parameter space allowing intermediate investments (0 < *c*\* < *r*). In the limit of zero within-patch growth, the allocation decision becomes a ‘zero-sum’ game, with no allowance for intermediate investment:

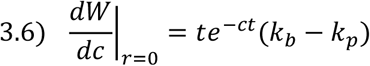

In this no growth scenario (*r* = 0), if biofilm cells have greater transmission value (*k_b_* > *k_p_*), then total biofilm investment is favored; if not, then total planktonic cell investment is favored, giving a strict trade-off defined by whichever fraction is preferentially transmitted between patches.

This simple zero-sum logic is intuitive, but fails significantly under more permissive growth conditions (i.e. increasing *r* and/or *t*; Figure 3 and Figure S2), where we see an intermediate level of colonization is favored over a relatively large portion of the parameter space. Despite large transmission advantages to biofilm cells, the intermediate colonization regime extends to the boundary of *k_p_* = 0 (black dashed line undefined for *r* > 0.04, Figure 3), such that even when planktonic cells have zero probability of founding a new patch, the vast majority of the population is still allocated to that fraction. This result follows from the dynamics of the within-patch model (Equations 2.1, 2.2 and Figure 2): when *k_p_* = 0, fitness is determined entirely by the size of the biofilm population, which is maximized at low colonization rates under conditions favoring growth due to the driving force of the planktonic growth engine.

### Biofilm as a bet hedge against environmental instability

In our evolutionary model (Figure 3), we assume that lineages can adapt their allocation decision making (*c*\*) in the context of constant transmission weightings *k_p_* and *k_b_*. However, the relative success of biofilm vs. planktonic cell propagules is likely to vary extensively in time as a function of unpredictable biotic and abiotic stresses (i.e. changing *k_p_* relative to *k_b_*). Despite reduced colonization rates leading to larger planktonic populations, sharply diminishing returns from further decreases in *c* (Figure 2E) suggest that the biofilm compartment has the potential to function as a cost-effective hedge against unpredictable selective events. At all growth rates, populations can exchange a small reduction in the size of the planktonic population for a massive increase in the biofilm population by raising the rate of colonization from a minimal value: returning to Equations 1.1 and 1.2, taking the ratio of the biofilm to planktonic growth rate yields the fraction 
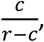
 which approaches infinity as *c* approaches *r*.

Maintaining a small biofilm presence is therefore relatively low-cost, even in the absence of environmental stresses that favor biofilm cells, suggesting that selection against low rates of biofilm production will be weak. To test this hypothesis, we performed a selection experiment, transferring 12 replicate populations of *P. aeruginosa* for 20 transfers (approximately 100 generations), allowing only planktonic or only biofilm cells to survive (Figure 4). While relative biofilm production rapidly decreased in the plankton passaged lines (*k_b_* = 0, *k_p_* = 1, red points in Figure 4A), declining from ~40% to ~13% of cells in biofilms after seven transfers (paired t-test, mean difference = 0.265, t = 18.7, df = 11, p = 1.1×10^−9^), biofilm production did not evolve to be any lower over the remaining 13 transfers of the experiment (Figure 4A, fraction biofilm from passages 7-20 best fit by an intercept model AIC = −580] vs. a linear model [AIC = −424]; also note in Figure 4C the biofilm OD holds roughly constant over the course of the experiment).

**Figure 4:**
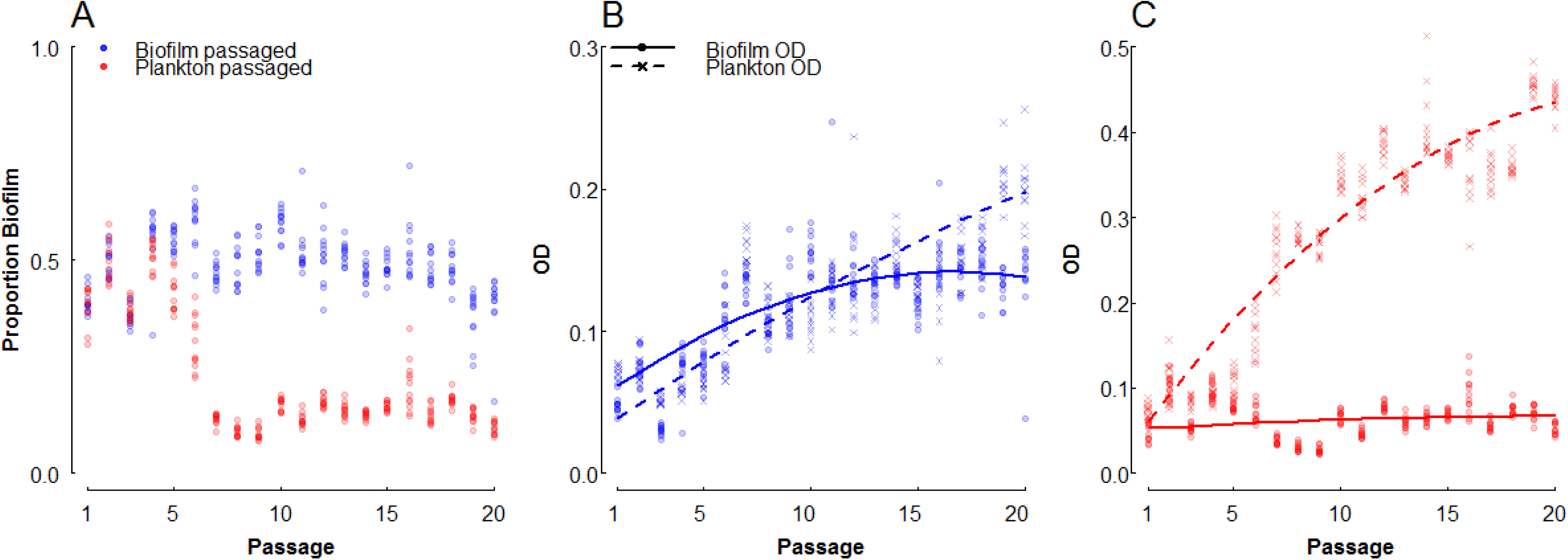
Selection for planktonic transmission fails to purge biofilm from experimental populations. *P. aeruginosa* (PAO1) was grown in 96-well plates containing a glass bead at room temperature, andevery 12 hours populations were fractionated, measured, and passaged to a new well. **A**, Proportion biofilm (defined as the ratio of the biofilm optical density (OD) to the sum of the ODs of the two fractions), **B**, fractionated ODs of B-selected (*k_b_* = 1, *k_p_* = 0), and **C**, fractionated ODs of P-selected (*k_b_* = 0, *k_p_*= 1) lines over the course of the passaging experiment. Points represent 12 independently evolving lineages per treatment, and curves from best-fit regression (adjusted R^2^ = 0.92, Table S1). All ODs are reported corrected for the OD of blank medium.

Interestingly, in the biofilm-only selection line (*k_b_* = 1, *k_p_* = 0; Figure 4A, blue points), the proportion of cells in biofilms did not increase over the course of the experiment, despite the total number of cells in the biofilm increasing by more than 100% (Figure 4B). Particularly, we see that the planktonic cell density increases steadily, while the biofilm fraction increases initially then stagnates after roughly 10 passages (Figure 4B, Table S1). This result is consistent with our models above, with planktonic growth driving biofilm accumulation, and experimentally demonstrates the utility of a mixed strategy under strict selection regimes.

The low cost of maintaining biofilm makes it well suited as a potential bet-hedging strategy, increasing the long-term geometric mean fitness in unpredictable environments by minimizing the variance in fitness through time. To test this prediction, we used our model framework to construct a simulated passaging regime in which growth and transmission parameters were subject to differing levels of variance. We assigned inoculum populations of planktonic cells a fixed colonization rate *c* (applied relative to *r*), which were then subjected to alternating periods of growth and transmission. Parameters *r*, *t*, *k_p_* and *k_b_* were drawn from normal distributions with fixed means µ, and differing variances σ^2^ between treatments. The values for *r*, *t* and *c* were used to solve Equations 2.1 and 2.2; proportions *k_p_* and *k_b_* of these new cells were then passaged to the next growth phase, as in Equation 3.1. The fitness function defined in equation 3.2 (effectively a reproductive number R0 (14)) was used as our fitness metric for these simulations, calculated as the ratio of founding cells in a given passage relative to the number of founding cells in the previous passage. Simulation results are displayed in Figure 5.

**Figure 5:**
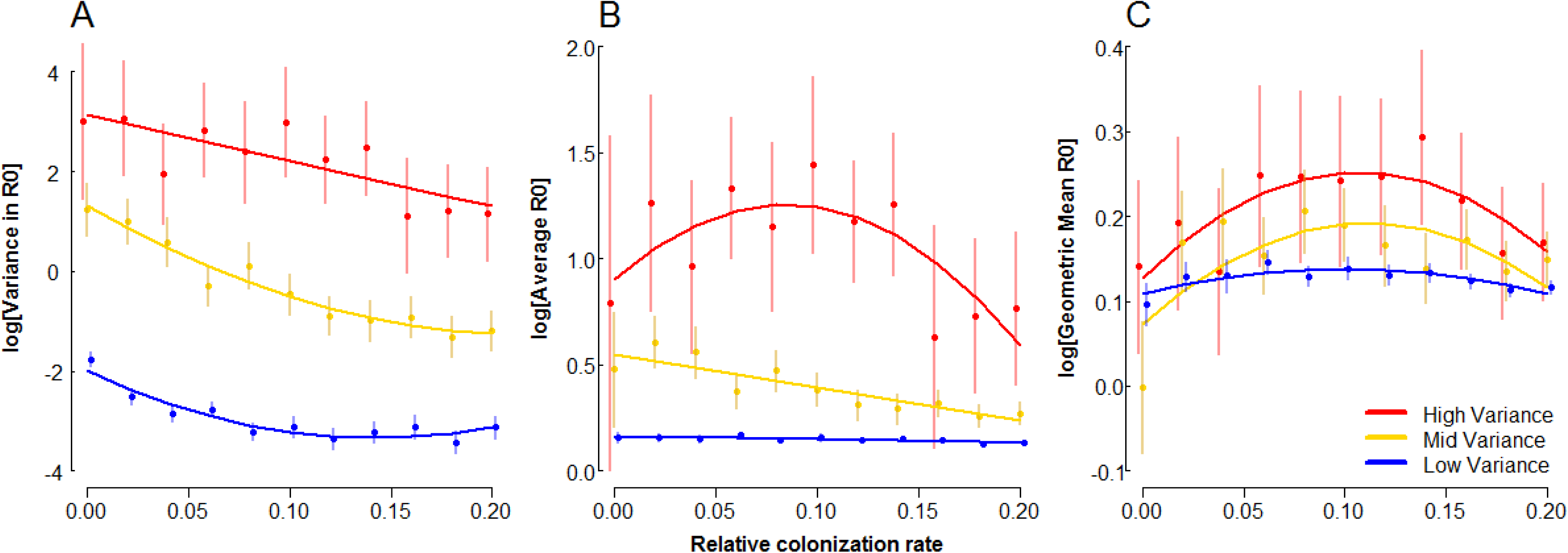
Biofilm production can act as a bet-hedging strategy. Log-scale plots of **A**, variance in R0, **B**,average R0, and **C**, geometric mean of R0, are plotted as a function of relative colonization rate (ratio of *c* to the expected value for *r*, see below) for lineages subject to passaging simulations with differentlevels of environmental instability. 100 replicate inocula of 5000 planktonic cells with the same fixed and µkb = 0.6. Under fixed conditions these parameters would favor *c*\* = 0.49. Variance treatments modified σ^2^ of the distributions from which *r*, *t*, *k_p_* and *k_b_* were drawn, with σ^2^low = µ/100, σ^2^mid = µ/9 andσ^2^high = µ/2.75. Parameters were restricted to logical values, with 0 ≤ *kb*, *kp* ≤ 1, *r* > 0, and *t* ≥ 1; this thresholding did not change the overall mean of any parameter by more than ±3%. R0 was calculated as the ratio of founding cells at a given passage relative to the founding cells of the previous passage. Points and error bars (offset to reduce overlap) represent means with 95% confidence intervals, and solid lines display best-fit regressions (Table S2).

For all treatments, the variance in R0 declines with increasing *c*, consistent with biofilm colonization functioning as a bet-hedging strategy for bacteria facing unpredictable selection (Figure 6A). For low and medium degrees of unpredictability, the average R0 declines monotonically with increasing *c*, reflecting the penalty imposed by environmental variance on fitness (Figure 6B). Interestingly, under the high variance treatment, a pronounced hump shape in average R0 appears (Table S2), indicating that under extremely unpredictable environments the trade-off between mean fitness and reduced variance in fitness breaks down and biofilm formation is generally beneficial (Figure 6B). The combination of direct fitness benefits and bet hedging effects lead to maxima in geometric mean R0 at intermediate colonization rates across all variance treatments (Figure 6C).

## DISCUSSION

In this work, we constructed a simple deterministic model of allocation between coupled biofilm and planktonic compartments within a growing bacterial population. Within the biofilm, cell division limited by geometry and nutrient diffusion rendered its effects inconsequential to the dynamics of the population as a whole, relative to the exponentially expanding pool of planktonic cells. Under inhospitable conditions, in which the population experienced restricted growth and/or frequent disruption, trade-offs between the biofilm and the planktonic compartment forced lineages to specialize in whichever fraction was favored to transmit between patches. When conditions become more permissive, the lineage is able to leverage exponential planktonic growth to maintain robust populations in both compartments, and at a fraction of the cellular cost of direct biofilm allocation (i.e. at reduced colonization rates *c*); in general, such cost saving measures are likely to be favored in any cooperative trait during periods of growth (15). Because maintenance of a biofilm comes at little cost under conditions favoring planktonic growth, biofilm is able to function as a robust and cost-effective hedge against unpredictable environmental change. Conversely, the planktonic fraction is a useful amplifier of biofilm cells even when biofilm is the transmissible propagule – this growth/transmission division of labor is more obvious for strictly non-growing phenotypes (e.g. spores), but the same logic holds for slow growth in biofilms as well.

We focus primarily on biofilm-planktonic cell populations as a model of survival-fecundity alternate states, and therefore use the language of colonization (of biofilms by planktonic cells) and dispersal (from the biofilm) to represent the switching processes between the two phenotypes. However, the model logic applies to other classes of resistant cells as well, such as persisters and spores. Indeed, there are instances where biofilm formation functions as a prerequisite or amplifying step in the formation of other types of resistant cells: persister cells are often enriched in biofilms (8, 16–19), and biofilm formation is a prerequisite step in fruiting body formation (the preferential site of sporulation) in *Bacillus subtilis* (20, 21); it would be interesting to investigate how investment would be optimized with multiple survival phenotypes available in both simultaneous and sequential contexts.

Given the costs inherent to cellular investment into a growth-limited state, one may expect lineages to evolve the ability to efficiently switch between biofilm and planktonic phenotypes, thereby optimizing fitness by reducing lag times and minimizing unnecessary mortality in the event of environmental disturbance, and indeed such systems appear to be abundant in nature (22–27). However, we note that environmental sensing in this case would not supplant the need to maintain some level of biofilm as a hedge (Figure 5, Figure 6), but rather to enhance the efficiency of such maintenance; in environments where catastrophic disturbances occur even at very low frequency, lineages that maintain biofilm regardless will still have better chances of survival. One would therefore predict biofilm to be completely lost in only the most constant environments (Figure 5).

Phenotypic regulation to further optimize allocation between biofilm and planktonic lifestyles would also help expedite the evolution of rudimentary life cycles at the population level, alternating between a growing ‘soma’ and dispersive ‘propagules’ with distinct demographies associated with each phase. Under the formalism presented here, either the biofilm or planktonic compartments alone, or some combination thereof, may serve as dispersing propagules. The historical archetype has generally held that the biofilm functions as the soma, with motile planktonic cells as dispersive propagules (28–30).

More recently Hammerschmidt et al. (31) found that alternating selection on dispersive and biofilm phenotypes in *Pseudomonas fluorescens* leads to the evolution of a lifestyle in which cooperative biofilm cells producing shared adhesive molecules form a pellicle that functions as the growing soma, and planktonic non-adhesive cheats are co-opted as dispersive propagules, thereby dividing labor between the two cellular fractions and increasing the overall fitness of the lineage. Our results indicate that the opposite cycle (biofilm cells as propagules, planktonic cells as soma) could also be viable, as the population can still reap the benefits of dividing labor between specialized cellular fractions. Indeed, where individual patches are permissive to growth, but transmission between patches is exceedingly harsh (e.g. wind-or animal-dispersal), dispersal via biofilm aggregates and growth within a planktonic ‘soma’ would offer the greatest advantage, as the ‘soma’ would accumulate biomass rapidly, and dispersal propagules would enjoy increased survival at little reproductive cost given the hostile transmission conditions. For example, biofilm formation and other survival phenotypes are likely important to successful transmission via fomites, upon which bacteria can remain viable for months (32). Dispersal in physically linked groups (i.e. budding dispersal, (33)) may also help maintain cooperative traits during dispersal, thereby potentially accelerating colonization when a new patch is reached. The biofilm ‘streamers’ observed by Drescher et al. (34) may be another example of this mode of transmission, where flow rates are such that the biofilm forms physical bridges to allow colonization of vacant surfaces in a topographically complex environment, as full detachment would prevent recolonization of adjacent surfaces due to extreme shear forces.

Taken together, our results highlight the evolutionary significance of within-population phenotypic heterogeneity and its consequences for survival and fecundity in mixed transmission environments. By optimizing the switching rate between robust and fecund specialists (here, the colonization rate from the planktonic to biofilm fractions, though we note that other mechanisms could lead to equivalent outcomes, such as the steady-state frequencies of genotypes arising from within-population diversification, as in (35, 36)), lineages were able to maximize fitness and transmission across a wide range of environments, as well as enhance survival in the face of catastrophic changes within the environment. The rate of phenotypic switching is therefore an essential parameter upon which selection may act when multiple phenotypes can persist within lineages.

## METHODS

### Passaging experiment

A mid-exponential phase culture of *P. aeruginosa* PA01 was used to inoculate 200 uL LB and one 3 mm sterile glass bead in each of 24 wells in a 96 well plate at an OD of 0.05. Plates were sealed with Aeraseal tape and grown statically at 24 °C in a humidified chamber. Every 12 hours (growth conditions were chosen to prevent entry into stationary phase, where multiple regulatory systems that modulate biofilm and growth behaviors are engaged), biofilm allocation was measured by removing and measuring the OD of the liquid phase, then washing, resuspending and measuring the density of the attached biofilm as above in 200 uL fresh LB. 12 lines had only the planktonic cells passaged, while the other 12 had only biofilm cells passaged; in each case, cells were diluted to an inoculum OD of 0.05, and 20 passages were performed.

### Statistics and mathematical analysis

Agent based simulations were performed using iDynoMiCs (13), and analytical analyses were performed using Mathematica. Numerical modelling (37) and statistics were performed in R (38), unless otherwise noted.

## ACKNOWELDGEMENTS

We thank R. Parthasarathy, B. Bohannan, and members of the Brown laboratory for helpful comments. The authors declare no conflicts of interest.

### FUNDING STATEMENT

NL was supported by a scholarship from the School of Biological Sciences, University of Edinburgh. LM was supported by Wellcome Trust grant #WT095831. WCR was supported by the Packard Foundation. LM and SPB were supported by HSFP grant #RGP0011/2014. The funders had no role in study design, data collection and interpretation, or the decision to submit the work for publication.

